# Siderophores provoke extracellular superoxide production by carbon-starving *Arthrobacter* strains when carbon sources recover

**DOI:** 10.1101/2020.03.20.000083

**Authors:** Xue Ning, Jinsong Liang, Yujie Men, Yangyang Chang, Yaohui Bai, Huijuan Liu, Aijie Wang, Tong Zhang, Jiuhui Qu

## Abstract

Superoxide and other reactive oxygen species (ROS) in the environment shape microbial communities^1^ and drive transformation of metals^2,3^ and inorganic/organic matter^4,5^. Taxonomically diverse bacteria and phytoplankton can produce extracellular superoxide during laboratory cultivation^6-11^. Understanding the physiological reasons for extracellular superoxide production by aerobes in the environment is a crucial question yet not fully solved. Here, we showed that iron-starving *Arthrobacter* sp. QXT-31 (referred to as *A.* QXT-31 hereafter) secreted a type of siderophore (deferoxamine, DFO), which provoked extracellular superoxide production by carbon-starving *A.* QXT-31 when carbon sources were recovered. Several other siderophores also demonstrated similar effects. RNA-Seq data hinted that DFO stripped iron from iron-bearing proteins in the electron transfer chain (ETC) of metabolically active *A.* QXT-31, resulting in electron leakage from the electron-rich (resulting from carbons source metabolism) ETC and superoxide production. Considering that most aerobes secrete siderophore(s)^12^ and often suffer from carbon starvation in the environment, certain aerobes are expected to produce extracellular superoxide when carbon source(s) recover/fluctuate, thus influencing the microbial community and cycling of many elements. In addition, an artificial iron-chelator (diethylenetriamine pentaacetic acid, DTPA) was widely used in microbial superoxide quantification. Our results showed that DTPA provoked superoxide production by *A*. QXT-31 and highlighted its potential interference in microbial superoxide quantification.

---

Reactive oxygen species (ROS), such as superoxide, are widely produced by aerobes and phytoplankton^6-11^. Biogenic ROS participate in interspecies signaling and microbial community shaping^1^ and drive the transformation of metals^2,3^ and organic matter^4,5^ in the environment. Single-electron reduction of oxygen produces superoxide, which can be further biologically/abiologically reduced to hydrogen peroxide (H_2_O_2_) and hydroxyl radical (HO^•^), hence superoxide production is of crucial importance. Although extracellular superoxide can be produced during normal aerobic metabolism of phylogenetically diverse aerobes^7^, the physiological reasons involved are far from clear. A better understanding of the physiological reasons for microbial ROS production would help to identify the influencing factors for microbial ROS production in natural ecosystems, and for the application of microbial ROS in bioremediation.

Iron in aerobic environments is commonly oxidized by oxygen into the ferric form (Fe(III)). To overcome the low bio-availability of Fe(III), most aerobes synthesize and secrete at least one type of siderophore^12^, which are widespread in the environment, to transport Fe(III) into cells. Here, we report on the novel role of siderophores in facilitating extracellular superoxide production by carbon-starving *Arthrobacter* strains when carbon sources recover. We observed that extracellular superoxide was not detected (using our earlier protocol^13^ based on the reaction between superoxide and superoxide-specific chemiluminescence (CL) probe MCLA^8^) in 48-h *A.* QXT-31 cultures grown in liquid mineral salt medium (MSM; containing 250 mg•L^-1^ glucose as the sole carbon source and 1.78 μM FeCl_3_; Supplementary Text 1), whereas glucose re-supplementation (50 mg/L) in 48-h *A.* QXT-31 cultures induced substantial superoxide production within approximate 30 min (Fig. 1a). Similar phenomena were also observed in the cultures of two other *Arthrobacter* strains (i.e., *A. cupressi* and *A. humicola;* purchased from China General Microbiological Culture Collection Center (CGMCC)) grown in modified peptone-yeast extract-glucose (mPYG) medium^14^ (Supplementary Fig. 2). After glucose was re-supplemented in the 48-h *A.* QXT-31 culture, the glucose concentration consistently decreased at first and then remained at a constant level (∼10 mg/L); superoxide CL signal intensity initially increased and peaked when the glucose concentration reached the constant value, and then persistently declined (Fig. 1a). Glucose re-supplementation only provoked superoxide production in *A.* QXT-31 cultured for more than 36 h (Fig. 1b), and maximal production in *A.* QXT-31 culture peaked on the sixth day and then declined (Fig. 1b). These observations of *A*. QXT-31 raised two questions: 1) what is the role of glucose in provoking superoxide production in old (≥36 h) *A.* QXT-31 cultures; and 2) why did glucose re-supplementation only trigger substantial superoxide production in ≥36-h *A.* QXT-31 cultures.

**Fig. 1.**
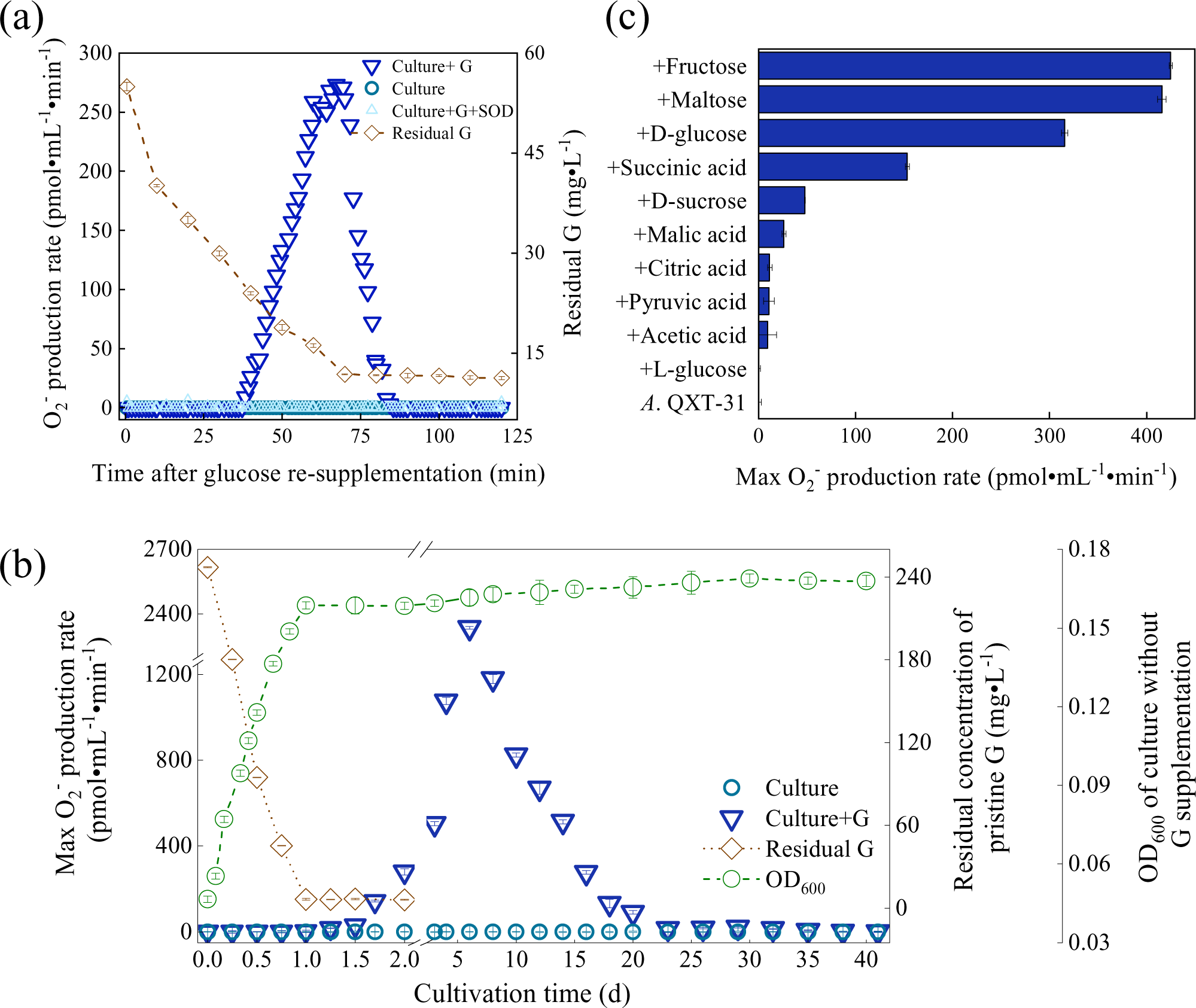
Superoxide production in *A*. QXT-31 culture re-supplemented with different carbon sources. (a) Superoxide CL signal intensity (only one of three biological replicates is shown here, with the other two shown in Supplementary Fig. 1) and residual glucose concentration (n = 3) in 48-h *A.* QXT-31 culture re-supplemented with/without sterile glucose (50 mg•L^-1^). Superoxide was monitored on a microplate reader after 180 μL of culture was added to each prepared microplate well (reagents added in advance; n = 3). Wells with SOD (120 kU•L^-1^) added were regarded as superoxide-free controls. (b) Maximum superoxide production rate, residual glucose concentration, and optical density (OD_600_) (n = 3) in *A.* QXT-31 cultures with/without sterile glucose re-supplementation (50 mg•L^-1^) during 40-d cultivation. (c) Maximum superoxide production rate in 48-h *A.* QXT-31 cultures with/without re-supplementation of fructose, maltose, D-glucose, succinic acid, D-sucrose, malic acid, citric acid, pyruvic acid, acetic acid, or L-glucose (n = 3). Carbon source re-supplementation in *A.* QXT-31 culture was consistent with carbon source upon which *A*. QXT-31 previously lived, with the exception of L-glucose treatment, where L-glucose was supplemented to *A*. QXT-31 culture pre-grown in D-glucose. Each carbon source was re-supplemented at the same concentration of 50 mg•L^-1^. Data are means ± average deviation of three replicates. G: Glucose.

Our results indicated that aerobic metabolism of glucose was required for superoxide production: i.e., 1) superoxide CL signal intensity in 48-h *A.* QXT-31 culture re-supplemented with sterile glucose (50 mg•L^-1^) stopped increasing right after glucose consumption ceased (Fig. 1a); 2) non-metabolizable L-isomer of glucose (L-glucose), instead of metabolizable glucose (D-isomer of glucose; D-glucose), was unable to induce production of extracellular superoxide (Fig. 1c); and 3) when *A.* QXT-31 was grown in MSM with other metabolizable carbon sources, carbon source re-supplementation in the culture induced superoxide production (Fig. 1c). The citric acid cycle, where electrons are generated during the breakdown of organic fuel molecules, is a primary metabolic process of these carbon sources. Thus, we speculated that glucose, and other carbon sources, probably provoked superoxide production by generating electrons during aerobic metabolism.

However, glucose re-supplementation did not induce substantial extracellular superoxide production in young (<36 h) *A.* QXT-31 cultures (Fig 1b). Rather, cells and extracellular substances in old (≥36 h) *A.* QXT-31 cultures appeared to favor superoxide production. As expected, when 24-h *A.* QXT-31 cells, deposited by centrifugation (718 *g*, 30 °C, 10 min), were suspended with the cell-free filtrate (CFF) of 48-h *A.* QXT-31 culture, glucose re-supplementation induced superoxide production in the suspension (Fig. 2a). These results demonstrated that extracellular substance(s) in 48-h *A.* QXT-31 culture induced 24-h cells to produce superoxide after glucose re-supplementation. The <3 kDa CFF fraction in 48-h *A.* QXT-31 culture was the only fraction capable of triggering superoxide production by 24-h cells (Fig. 2b). Chromatographic analysis of fresh MSM and <3 kDa CFF fractions of 24- and 48-h *A.* QXT-31 cultures showed that a conspicuous peak signal was only observed in the 24- and 48-h CFF, with the peak in 48-h CFF approximately double that in 24-h CFF (Supplementary Fig. 3). The exclusive peak in CFF was then identified by electrospray ionization tandem mass spectrometry (ESI-MS/MS). Its primary and secondary mass spectra shared a predominant m/z peak and molecular ion fragmentation with that of deferoxamine (DFO) (Supplementary Fig. 4&5), a common type of microbial siderophore. The DFO standard (deferoxamine mesylate salt; European Pharmacopoeia Reference Standard) shared a similar retention time (12.51 min) as the exclusive peak in CFF (12.49 min) (Supplementary Fig. 3). The addition of 2.0 μM of DFO (approximate to the difference in DFO concentration between 24- and 48-h cultures) into glucose-re-supplemented 24-, 36-, and 48-h *A.* QXT-31 cultures triggered/enhanced superoxide production (Fig. 2c). A siderophore biosynthesis gene (locus tag BWQ92_RS08305) was predicted in the *A*. QXT-31 genome using the NCBI Prokaryotic Genome Annotation Pipeline^15^ (website provided in Methods section), indicating that *A*. QXT-31 is capable of synthesizing at least one type of siderophore. Our results strongly demonstrated that *A*. QXT-31 synthesized and secreted DFO during cultivation (probably as a responding to iron starvation), which accumulated extracellularly and provoked carbon-starving cells to produce superoxide when the utilizable carbon source was recovered.

**Fig. 2.**
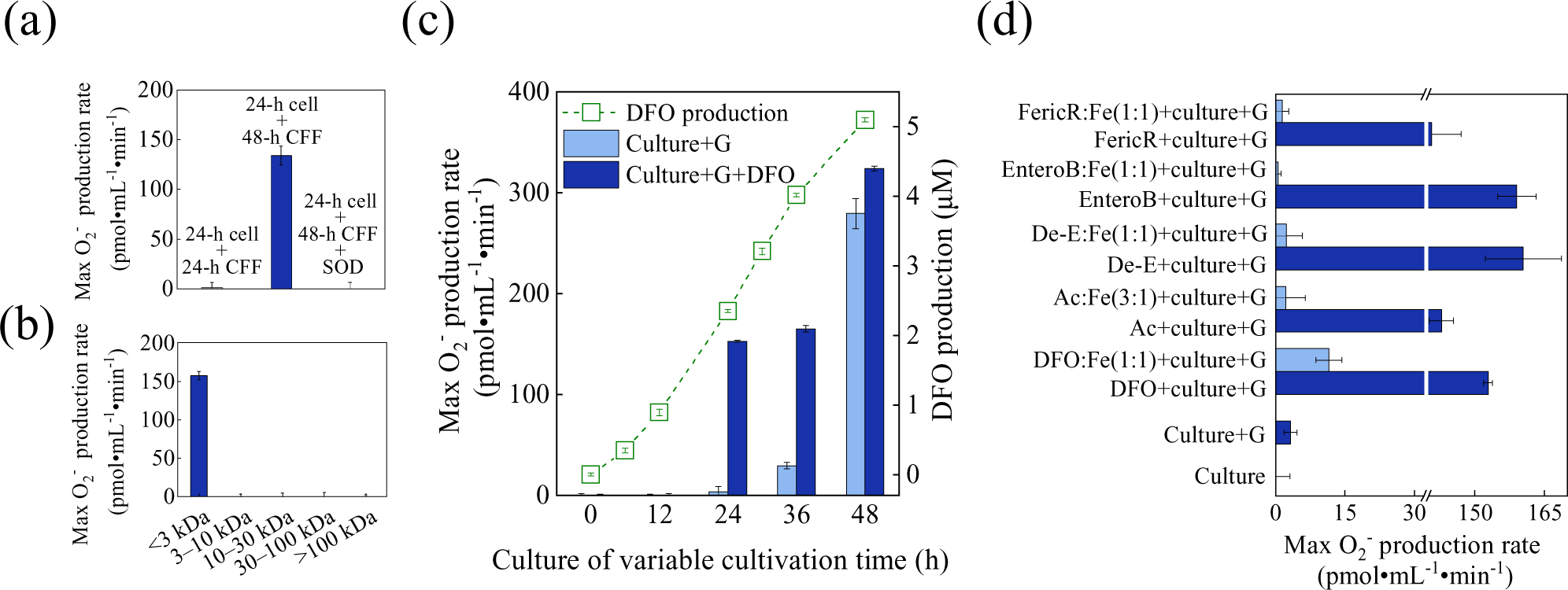
DFO in CFF of *A*. QXT-31 culture provoked superoxide production. (a) Maximum superoxide production rate in 24-h *A*. QXT-31 culture and 24-h *A*. QXT-31 cell suspension in CFF of 48-h *A*. QXT-31 culture. SOD (120 kU•L^-1^) was added to generate superoxide-free controls. Superoxide CL data were collected on a microplate reader after cultures/suspensions were mixed with 50 mg•L^-1^ glucose in microplate wells (n = 3). (b) Maximum superoxide production rate in 24-h *A*. QXT-31 cell suspension (with 50 mg•L^-1^ glucose re-supplementation) in 48-h CFF fractions of <3 kDa, 3–10 kDa, 10–30 kDa, 30–100 kDa, and >100 kDa. (c) DFO production in *A*. QXT-31 culture, and maximum superoxide production rates in cultures (with 50 mg•L^-1^ glucose re-supplementation) with/without DFO addition (2 μM; approximate to DFO concentration in 24-h culture). (d) Maximum superoxide production rate in 24-h *A*. QXT-31 cultures (with/without 50 mg•L^-1^ glucose re-supplementation) with/without each free/Fe(III)-preincubated siderophore. FericR: ferrichrome; EnteroB: enterobactin; De-E: deferrioxamine E; Ac: Acetohydroxamic acid. Data are means ± average deviation of three replicates. G: Glucose.

The affinity of DFO to Fe(III) under physiological conditions is much greater than that of the common artificial metal-chelator, ethylene diamine tetraacetic acid (EDTA)^16^. Hence, it was hypothesized that DFO promotes superoxide production by exploiting its high affinity to Fe(III). This hypothesis was supported by the following experimental results: 1) Each of the four other types of iron-free siderophore (e.g., acetohydroxamic acid, deferrioxamine E, enterobactin, and ferrichrome) also triggered superoxide production in glucose-re-supplemented 24-h *A.* QXT-31 culture (Fig. 2d); however, 2) superoxide production was attenuated by preincubating the siderophores with Fe(III) (Fig. 2d).

The underlying mechanism involved in the facilitation of superoxide production by DFO was further explored using RNA-Seq. Their high affinity for Fe(III) enables certain siderophores to strip iron from iron-bearing proteins^17,18^, and thus diminish their activities. Accordingly, DFO was suspected to strip iron from iron-bearing proteins of *A*. QXT-31 and thus inactivate these proteins. RNA-Seq analysis showed that DFO supplementation (2.0 μM; approximate to DFO concentration in 24-h culture) up-regulated the transcriptional level of genes encoding iron-related and iron-bearing proteins (including Fe-S cluster assembly proteins, NADH dehydrogenase, ubiquinol-cytochrome C reductase, cytochrome B, and ferredoxin) in *A.* QXT-31 cultures (without carbon source supplementation) at different ages (12, 24, 36, and 48 h). The highest up-regulation was observed in the 48-h culture (Fig. 3a and Supplementary Fig. 6a). Transcriptional up-regulations of the same genes were also observed in 36-h *A*. QXT-31 cells treated by the other siderophores (2 µM), compared to cells without siderophore treatment (Fig. 3b). An increase in Fe-S cluster synthesis could be a response to Fe-S cluster damage in proteins^19^. Hence, the above results suggest that DFO stripped iron from these iron-bearing proteins and thus impaired their activities. ETC in bacterial plasmalemma are functionally similar to that of eukaryotic mitochondria. In addition, mitochondrial H_2_O_2_ production (biogenic H_2_O_2_ is easily transformed from superoxide by the superoxide-producing cells themselves and is commonly used for indirect quantification of superoxide) can be promoted by electron transfer inhibitors by disturbing the electron transfer process^20,21^. The transcriptional up-regulation of genes encoding NADH dehydrogenase (ETC complex I) and ubiquinol-cytochrome C reductase (ETC complex III) was observed after DFO addition (Fig. 3a), indicating that DFO caused the impairment/disfunction of the two complexes in the ETC of *A*. QXT-31 cells. Considering perturbations of ETC functions of eukaryotic cells promoted superoxide production during the consumption of NADH^20,21^, the above results suggest that DFO-induced disfunction of electron-rich (resulting from carbons source metabolism) ETC complexes I and III was a probable reason for superoxide production by metabolically active *A*. QXT-31.

**Fig. 3.**
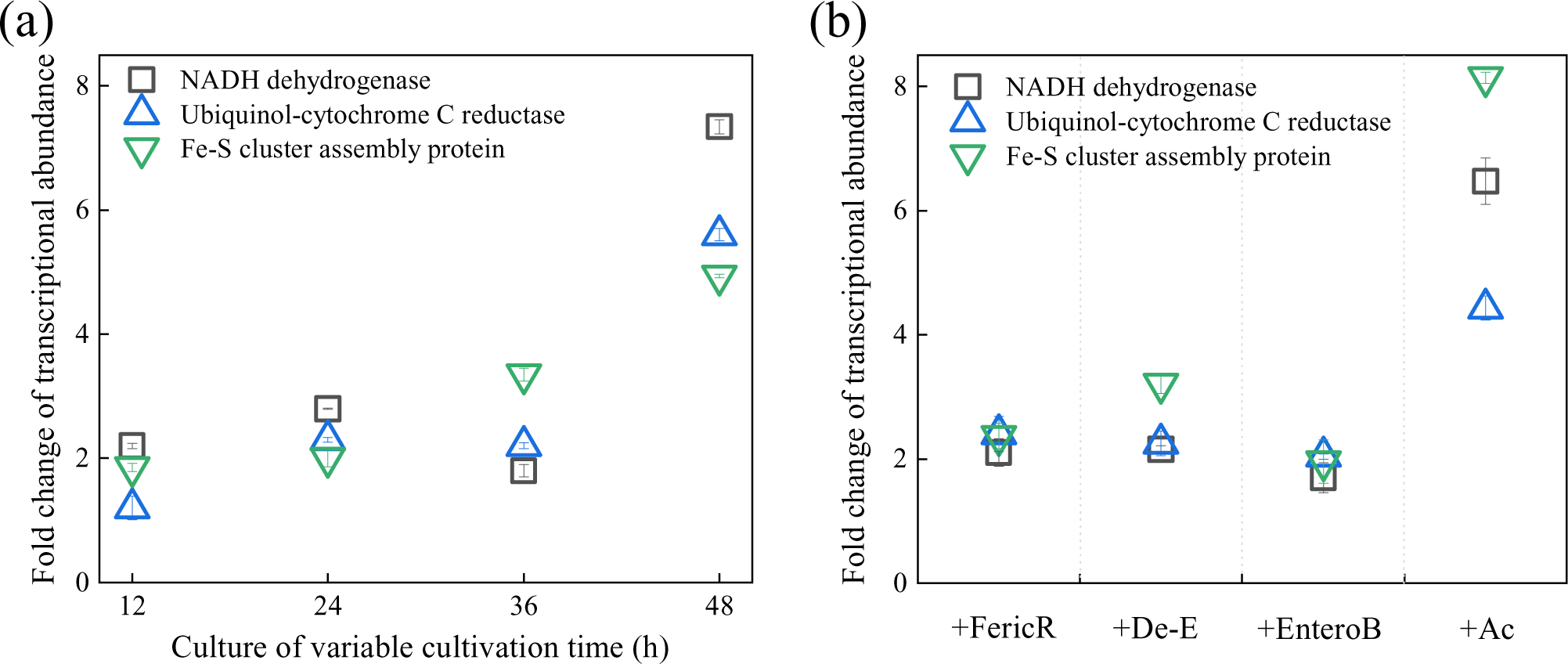
Siderophores caused transcriptional up-regulation of genes encoding iron-related and iron-bearing proteins. (a) Changes in the transcriptional abundance of genes encoding iron-related and iron-bearing proteins after 2.0 μM sterile DFO was added to *A.* QXT-31 cultures. After DFO addition, cells in 0.5 mL of culture were harvested by centrifugation (10□000 *g*, 4 °C, 3 min) at 0.5 h, 1 h, 1.5 h, and 2 h, with four cell samples collected at four time points then mixed well as an RNA-Seq sample. Cells in *A*. QXT-31 cultures without DFO addition were used as controls. (b) Changes in the transcriptional abundance of genes encoding iron-related and iron-bearing proteins in 36-h *A.* QXT-31 culture after supplementation with (2 μM) acetohydroxamic acid (Ac), deferrioxamine E (De-E), enterobactin (EnteroB), and ferrichrome (FericR). RNA-Seq samples were prepared according to that of DFO. Data are means ± average deviation of two biological replicates.

The above findings prompted a rethink of the methodology of cellular superoxide quantification, where a metal-chelator, DTPA, was widely used^7-10,22,23^. DTPA was initially exploited in a superoxide producing system (xanthine-xanthine oxidase system; used to generate superoxide at an expected rate) to maintain superoxide signals by suppressing interference from metal ions^23,24^. However, as a Fe(III)-chelator, DTPA may enhance/provoke cellular superoxide production, resulting in overestimated or false-positive results when estimating cellular superoxide production rates under physiological conditions. Our data showed that DTPA induced *A.* QXT-31 to produce more extracellular superoxide, even at concentrations (5–10 μM) (Fig. 4a) lower than used in previous research (40–170 µM)^7-10,22^. Similar phenomena were also observed when DTPA was replaced with DFO, EDTA, and acetohydroxamic acid (Supplementary Fig. 8). Increase in superoxide production in the DTPA-added *A.* QXT-31 culture may result from two possibilities: 1) cells were altered by DTPA to become superoxide producer, and 2) extracellular superoxide scavenger in *A.* QXT-31 culture was suppressed by DTPA, leaving preexisting superoxide detected. Compared to 24-h *A*. QXT-31 culture re-supplemented with glucose (25 mg/L), DTPA-treated (10 μM; for 10 min) 24-h *A*. QXT-31 cells produced markedly more extracellular superoxide after resuspension in the glucose-re-supplemented (25 mg/L) raw CFF of 24-h *A*. QXT-31 culture (Fig. 4b), indicating that DTPA acted on and provoked *A*. QXT-31 cells to produce superoxide.

**Fig. 4.**
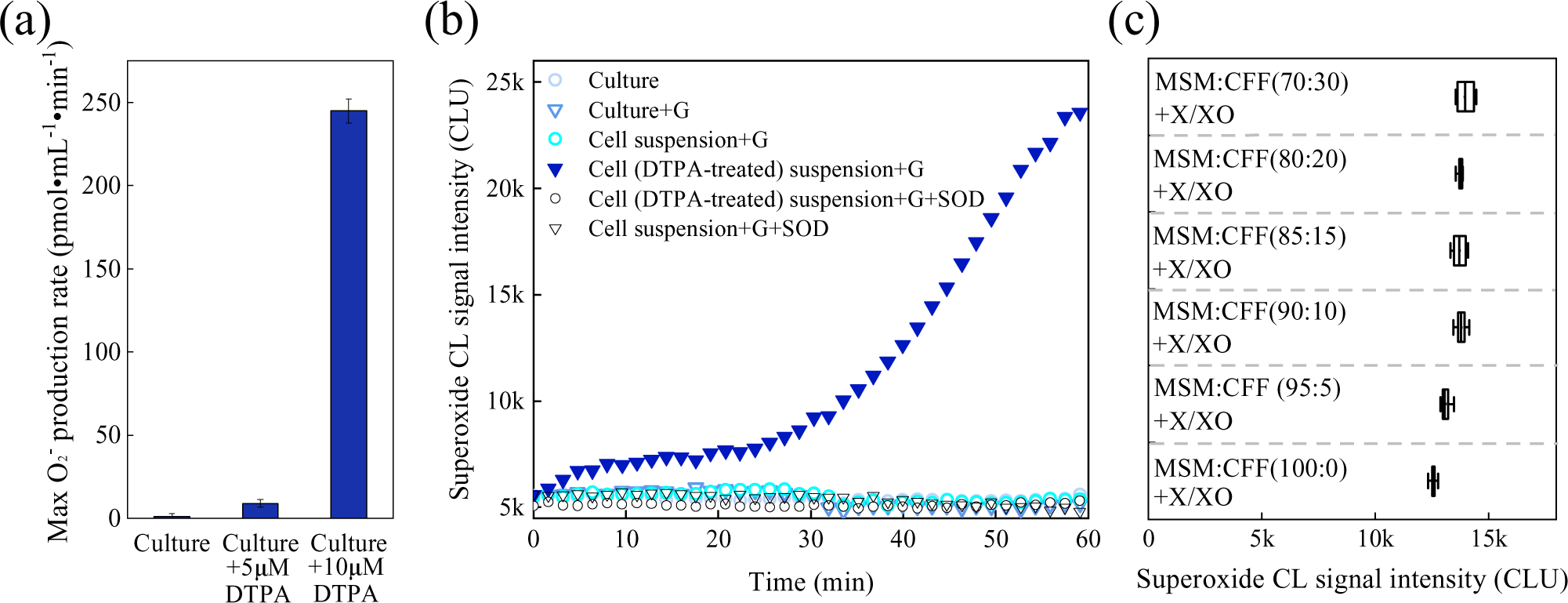
DTPA provoked *A*. QXT-31 cells to produce superoxide. (a) Maximum superoxide production rate in glucose-re-supplemented (50 mg•L^-1^) 24-h *A*. QXT-31 cultures with/without the addition of DTPA. (b) Superoxide CL signal intensity in glucose-re-supplemented (50 mg•L^-1^) 24-h *A*. QXT-31 cultures and suspensions (in CFF of 24-h culture) of DTPA-treated 24-h *A*. QXT-31 cells (only one of three biological replicates is shown here, with the other two shown in Supplementary Fig. 6). 24-h *A*. QXT-31 culture was supplemented with/without 10 μM sterile DTPA, followed by shaking at 170 rpm at 30 °C in an oscillating incubator for 10 min. Cells were collected by centrifugation (718 *g*, 30 °C, 10 min), and then suspended in CFF of 24-h culture (DTPA free). SOD (120 kU•L^-1^) was added to generate superoxide-free controls. (c) Superoxide CL signal intensity range in MSM with/without CFF of 24-h culture. MSM was first mixed with CFF to generate mixtures with variable volume ratios (100:0, 95:5, 90:10, 85:15, 80:20, and 70:30 MSM:CFF), then the mixtures were added to microplate wells preloaded with superoxide-producing reagents (250 μM xanthine (X) and 200 mU•L^-1^ xanthine oxidase (XO)) and MCLA (3.125 μM) for immediate determination in a microplate reader. Superoxide CL signal intensity was collected within 3 min (10 measurements for each treatment, n = 10) and compared. G: Glucose.

The CL signal intensity of superoxide produced by xanthine (X) and xanthine oxidase (XO) (250 μM X and 200 mU•L^-1^ XO) in modified MSM (without trace heavy metals or cofactors; with 50 mg•L^-1^ glucose) was not suppressed in the presence of CFF from 24-h culture (Fig. 4c), suggesting the absence of a superoxide scavenger in the CFF (Fig. 4c). Hence, these results indicate that the DTPA-induced superoxide increase in *A*. QXT-31 culture was totally attributable to the effect of DTPA on *A*. QXT-31 cells. Thus, DTPA and other metal chelators should be used with caution, as they may exert a positive influence on superoxide production in certain microbes and interfere cellular superoxide detection/quantification.

The finding also shed light on the important role of siderophores in microbial ecology. Aerobes produce and secrete over 500 different types of siderophores^25^, which accumulate in the environment due to their good stability. Hydroxamate-type siderophores in soil have been reported as high as 10 µM^26^. In addition, aerobes often suffer from carbon starvation in the environment^27^. When a small quantity of carbon (as low as 15 mg/L of glucose for *A*. QXT-31 (Supplementary Fig. 9)) becomes available for these Fe- and carbon-starving aerobes, some species (such as *Arthrobacter* species) of microflora can produce extracellular superoxide. As superoxide and other ROS are toxic to cells, the neighbors of extracellular superoxide producers may be suppressed, and superoxide producers, which should be resistant to ROS, will likely succeed in carbon source competition. Hence, some aerobes may change the microbial community by producing extracellular superoxide when carbon source levels fluctuate. In addition, superoxide produced by Fe- and carbon-starving aerobes during carbon source fluctuation may also accelerate the transformation of metals and inorganic/organic matter in the environment.

## Methods

### Bacterial strain and cultivation

We used an aerobic gram-positive strain of *Arthrobacter* sp. QXT-31 (referred as *A.* QXT-31) isolated from surface soil obtained from a manganese mine in Hunan Province, China^14^. The *A.* QXT-31 strain was deposited in the CGMCC (CGMCC number 6631). For experimentation, *A.* QXT-31 was grown in mineral salt medium (MSM, Supplementary Text 1). For each cultivation in liquid medium, an agar-plate colony was transferred into 30 mL of liquid culture for 48-h cultivation in the dark at 170 rpm and 30 °C. After subculturing for one generation, the bacterial culture grown for 24 h was used as an inoculum (3% inoculation proportion, v:v). Both *Arthrobacter cupressi* (CGMCC number 1.10783) and *Arthrobacter humicola* (CGMCC number 1.15654) were grown in modified peptone-yeast extract-glucose (mPYG) medium^14^. Cell growth was estimated using optical density at 600 nm (OD_600_) with the bacterial suspension, monitored by a Spark™ 10M microplate reader (Tecan, Switzerland).

### Extracellular superoxide production assay

A superoxide-specific chemiluminescent (CL) probe (MCLA, 2-methyl-6-(4-methoxyphenyl)-3,7-dihydroimidazo[1,2-a]pyrazin-3(7H)-one; TCI, Japan) was used for the extracellular superoxide assays^8^ using the microplate luminometer of the microplate reader. According to our previous study^13^, DTPA was excluded in the superoxide quantification system unless otherwise stated. Xanthine and xanthine oxidase from bovine milk (Sigma-Aldrich, USA) were added to the bacterial cultures to generate a calibration curve between the superoxide production rate and superoxide CL signal intensity for calibration of the bacterial superoxide production rate^8^. Superoxide dismutase (SOD) from erythrocytes of *Bos grunniens* (Gansu Yangtaihe Biotechnology Co., Ltd., China) was used to generate a superoxide-free control. Details are available in Supplementary Text 2.

### Glucose quantification

Glucose concentration in the bacterial cultures was measured using a glucose quantification kit (E1010, Applygen, China) based on the glucose oxidase/peroxidase method^28^ in accordance with the manufacturer’s instructions. A calibration curve between glucose concentration and optical density at 500 nm, which was determined using the microplate reader, was established to calibrate the glucose concentration in samples.

### Secretion fractionation and identification

Secretions of *A.* QXT-31 in MSM were fractionated based on molecular weight to explore the molecular weight range of the activated substance(s) facilitating superoxide production by *A.* QXT-31. Secretions in 48-h *A.* QXT-31 cultures were first centrifuged (Sigma 3-18KS, Germany) at 10□000 *g* and 4 °C for 5 min, with the resulting supernatant filtrated through a 0.22-μm sterile filter (Guangzhou Jet Bio-Filtration Co., Ltd, China) to prepare the cell-free filtrate (CFF). The CFF was then fractionated into five fractions (>100 kDa, 100–30 kDa, 30–10 kDa, 10–3 kDa, and <3 kDa) using Millipore ultrafiltration centrifugal filters with nominal molecular mass limits of 100 kDa, 30 kDa, 10 kDa, and 3 kDa. The 24-h *A.* QXT-31 cells were harvested by centrifugation (718 *g*, 30 °C, 10 min), and suspended in each of the five fractions. The suspensions (180 μL) were added to microplate wells preloaded with MCLA (3.125 μM), glucose (50 mg/L), and SOD (120 kU•L^-1^, only for controls), and superoxide CL signal intensity was detected immediately. An Ultimate 3000 ultra-high-performance liquid chromatography (UPLC) system, combined with a Q Exactive Plus mass spectrometer (Thermo Fisher Scientific, USA), was used to identify suspected substance(s) in the secretion fractions. Details are available in Supplementary Text 3.

### Preparation of Fe(III)-preincubated siderophores

Fe(III)-saturated siderophore solution was prepared by slowly adding freshly prepared Fe(III) solution (FeCl_3_·6H_2_O dissolved in deionized water) to the siderophore solution at a variable molar ratio (depending on Fe(III) complexing site number of siderophore molecules) so that the complexing site of each siderophore was saturated by Fe(III), leaving negligible uncomplexed Fe(III). The Fe(III)-preincubated siderophore solution was allowed to equilibrate for at least 1 h at room temperature before use.

### RNA extraction, sequencing, and transcriptome analysis

RNA-Seq was used to estimate transcriptional abundance of *A.* QXT-31 cells in variable conditions: 1) *A.* QXT-31 cells cultured for 12 h, 24 h, 36 h, and 48 h with/without exogenetic DFO (2 μM) treatment; and 2) 36-h *A.* QXT-31 cells treated by each of the four siderophores (2 μM; acetohydroxamic acid, ferrichrome, enterobactin, and deferrioxamine E). Each siderophore was added to the culture (5 mL) to react with cells for 2 h, and cells in 0.5 mL of the culture (5 mL) were then harvested by centrifugation (10□000 *g*, 4 °C, 3 min) at 0.5 h, 1 h, 1.5 h, and 2 h, with the four cell samples collected at the four timepoints mixed well as an RNA-Seq sample. TRNzol reagent (DP424, TIANGEN, China) was used for RNA extraction according to the manufacturer’s instructions, with modification of the cell lysis step, where cell pellets were pulverized by a pestle in liquid nitrogen^13^. Total RNA concentration, RNA integrity number (RIN), and RNA quality number (RQN) were evaluated using an Agilent 2100 Bioanalyzer (Santa Clara, USA). Samples with a RIN/RQN value above 8.0 were collected for further analysis. Extracted RNAs were kept at −80 °C before cDNA library construction. Details of RNA sequencing and transcriptome analysis are listed in Text S4. All raw sequences generated from RNA-Seq were deposited in the NCBI Sequence Read Archive database under accession number PRJNA607123.

The coding regions of all *A.* QXT-31 genes annotated by the NCBI Prokaryotic Genome Annotation Pipeline ^15^ (ftp://ftp.ncbi.nlm.nih.gov/genomes/all/GCF/001/969/265/GCF_001969265.1_ASM196926v1) were used as a reference for transcriptome analysis. Transcriptional abundance was estimated using a build-in script (*align_and_estimate_abundance.pl*) and normalized for cross-sample comparison using a build-in script (*abundance_estimates_to_matrix.pl*) with the Trinity platform (v2.8.5)^29^.

### Influence of DTPA on cellular superoxide production

The 24-h *A.* QXT-31 cultures were supplemented with/without 10 μM sterile DTPA and incubated at 170 rpm for 10 min at 30 °C, with 1 mL of each culture then centrifuged (718 *g*, 30 °C, 10 min) for cell deposition. Cells were suspended in 1 mL of CFF (DTPA free) from the 24-h *A.* QXT-31 culture. The suspensions (180 μL) were added to microplate wells preloaded with MCLA (3.125 μM), glucose (50 mg•L-1), and SOD (120 kU•L^-1^, only for controls), and the superoxide CL signal intensity was detected immediately.

The potential presence of an extracellular superoxide scavenger in 24-h *A*. QXT-31 cultures was explored. MSM (without heavy metals or cofactors; with 50 mg•L^-1^ glucose) was prepared and mixed with the CFF of 36-h *A.* QXT-31 culture at variable volume ratios (i.e., 100:0, 95:5, 90:10, 85:15, 80:20, and 70:30 MSM:CFF). The mixtures were added to microplate wells preloaded with superoxide-producing reagents (250 μM xanthine (X) and 200 mU•L^-1^ xanthine oxidase (XO)) and MCLA (3.125 μM) for immediate determination in the microplate reader. Superoxide CL signal intensity was collected within 3 min (10 measurements for each treatment) and compared.

### Chemicals

Chemicals used in this study included acetohydroxamic acid (Rhawn, China, >98% purity); ferrichrome (*Ustilago sphaerogena)* (Sigma-Aldrich, USA, >99%); enterobactin (*Escherichia coli*) (Sigma-Aldrich, USA, ≥98%); and deferrioxamine E (Abcam, UK, >95%).

## Acknowledgements

This study was supported by the National Natural Science Foundation of China (Funding No. 31700106, 51778603, and 51820105011). This research was also funded by the Hong Kong Scholars Program.

## Contributions

J.Q. and J.L. conceptualized the study. X.N. and J.L. performed experiments and data analysis. J.L. wrote the manuscript, with suggestions from Y.B., X.N., Y.M., T.Z., Y.C., H.L., and A.W..

