## Supplementary Information for "Siderophores provoke extracellular superoxide production by carbon-starving *Arthrobacter* strains when carbon sources recover"

**Supplementary Text 1**

**MSM composition and preparation process**

MSM composition: 500 mg•L^-1^ of MgSO_4_·7H_2_O, 60 mg•L^-1^ of CaCl_2_·2H_2_O, 4 g•L^-1^ of Na_2_HPO_4_·12H_2_O, 500 mg•L^-1^ of KH_2_PO_4_, 250 mg•L^-1^ of (NH_4_)_2_SO_4_, 0.5 mg•L^-1^ of FeCl_3_·6H_2_O, 0.5 mg•L^-1^ of Na_2_MoO_4_·2H_2_O, 0.5 mg•L^-1^ of CuCl_2_, 0.5 mg•L^-1^ of MnCl_2_·2H_2_O, 0.5 mg•L^-1^ of ZnSO_4_·7H_2_O, 0.05 mg•L^-1^ of NiCl_2_·6H_2_O, 0.001 mg•L^-1^ of vitamin B1, 0.001 mg•L^-1^ of vitamin B12, 0.001 mg•L^-1^ of biotin, 10 mM mg•L^-1^ of HEPES.

Preparation process: A mixed solution of MgSO_4_·7H_2_O and CaCl_2_·2H_2_O was autoclaved at 120 °C for 20 min. Stock solution (100 X) of Na_2_HPO_4_·12H_2_O, KH_2_PO_4_ and (NH_4_)_2_SO_4_ was sterilized by filtration before use. Metal stock solution (50 X) was prepared as follows: FeCl_3_·6H_2_O was first added and dissolved in pre-acidized (by hydrochloric acid, pH ~3.0) ultrapure water (18.3 MΩ) to suppress Fe(III) hydrolyzation, with each of the other metals (Na_2_MoO_4_·2H_2_O, CuCl_2_, MnCl_2_·2H_2_O, ZnSO_4_·7H_2_O, and NiCl_2_·6H_2_O) then added and totally dissolved before the addition of the following metal. The metal stock solution was sterilized by filtration before use. The cofactor stock solution (1 000 X) contained vitamin B1, vitamin B12, and biotin. Carbon sources and HEPES were sterilized by filtration before use.

**Supplementary Text 2**

**Extracellular superoxide quantification**

Extracellular superoxide was quantified as per Godrant et al.^1^ with some modifications. A superoxide-specific chemiluminescent (CL) probe, MCLA (2-methyl-6-(4-methoxyphenyl)-3, 7-dihydroimidazo [1, 2-a] pyrazin-3(7H)-one), which emits light at 460 nm when reacting with superoxide, was added to the culture suspension, and the CL signal was monitored by a Spark™ 10M microplate reader (Tecan, Switzerland). Reaction between xanthine (X) and xanthine oxidase (XO) generates superoxide. Thus, X and XO were added to the culture suspensions to generate a calibration curve between the superoxide production rate and CL signal density and thus quantify superoxide produced by cells. Stock solutions of 125 µM MCLA (TCI scientific, Japan), 5 mM xanthine (Sigma-Aldrich), 3 U•L^-1^ XO, and 3 kU•mL^-1^ modified (for higher stability) copper zinc superoxide dismutase (SOD) from erythrocytes of *Bos grunniens* (Gansu Yangtaihe Biotechnology Co., Ltd., China) were prepared in 18.2 MΩ cm^-1^ Milli-Q water. Superoxide production rates in the X/XO system were calculated according to the following equation^2^:

$$\frac{d[O_{2}^{-}]}{dt}=\frac{0.618}{{10}^{pK_{1}-pH}+1+{10}^{pH-pK_{2}}}=nmol/U XO/s$$

where p*Κ*_1_ = 6.6 (p*K*a of xanthine oxidase) and p*Κ*_2_ = 8.2 (p*K*a of xanthine). The pH in the culture system was controlled at 7.3.

The superoxide production rate in the culture suspensions was quantified as follows. In brief, 10 μL of xanthine ([X]_final_ = 250 μM) was first added to the bottom of 15 wells in a white 96-well plate. For the blank, 5 μL of SOD ([SOD]_final_ = 120 kU•L^-1^) was added to the walls of three wells. For the standards, 5 μL of XO secondary standard 1 ([XO_1_]_final_ = 2.5 mU•L^-1^), standard 2 ([XO_2_]_final_ = 175 mU•L^-1^), and standard 3 ([XO_3_]_final_ = 350 mU•L^-1^) were added to the walls of three wells, respectively. The separated reagent drops in each well were mixed with 180 μL of culture suspension, and the background CL signals in the 15 wells were first read prior to MCLA addition, after which 5 μL of MCLA ([MCLA]_final_ = 3.125 μM) solution maintained at room temperature in the dark was added to each of the 15 wells and mixed with the other reagents by a pipette. The CL signals in all 15 wells were read using an acquisition time of 1 s per well.

After the carbon source was supplemented to any culture, superoxide CL signal intensity was monitored on a microplate reader for about 120 min. The maximum value of superoxide CL signal intensity was used to calculate maximum superoxide production rate.

**Supplementary Text 3**

**MS/MS identification**

**1. Sample preparation.** Samples were filtered through a 0.22-μm pore filter. Acetonitrile was chosen as the precipitant. A sample and acetonitrile mixture (volume ratio of 1:4) was prepared in a 1.5-ml Eppendorf tube, and mixed using a vortexer (Vortex-Genie 2T, Scientific industries, USA). The solution was then centrifuged at 18 000 *g* for 10 min at 4 °C, and the sediment was removed. The solution without sediment was then transferred to a vacuum concentrator (CentriVap, Labconco, USA) and was not taken out until all solvents were evaporated. Solutes left in the 1.5-ml Eppendorf tube were finally dissolved in 50 μL of Milli-Q water.

**2. Chromatographic separation.** Sample separation was performed on an Ultimate 3000 ultra-high-performance liquid chromatography (UHPLC) system (Thermo Fisher Scientific, USA). A Waters ACQUITY UPLC HSS T3 column (2.1 mm × 100 mm × 1.8 μm) was selected for chromatographic separation, with mobile phase composition of methanol containing 0.1% (v/v) formic acid (A) and an aqueous solution containing 0.1% (v/v) formic acid (B). The sample (2 µL) was injected into the UPLC system. The mobile phase was delivered at a flow rate of 0.2 mL·min^-1^ with a gradient elution program of: 100% B at 0–2 min, 100–0% B at 2–20 min, 100% A at 20–25 min, 0–100% B at 25–25.1 min, and 100% B at 25.1–30 min.

**3. High resolution mass spectrometry.** High-resolution mass detection was carried out on a Q Exactive Plus mass spectrometer (Thermo Fisher Scientific, USA) combined with UHPLC through both negative electrospray ionization (ESI^-^) and positive electrospray ionization (ESI^+^) interface simultaneously. In ESI^-^ mode, the sheath gas and auxiliary gas flow rates were set at 30 and 10 Arb, respectively. The spray voltage was maintained at 2.80 kV. In ESI^+^ mode, the sheath gas and auxiliary gas flow rates were set at 35 and 10 Arb, respectively. The spray voltage was maintained at 3.50 kV. The stepped normalized collisional energies were 20, 40, and 60 eV. The capillary temperature was maintained at 320 °C. Full scan mass data were collected from 200 to 3 000 m/z with a resolution of 70 000 full width at half maximum (FWHM) in continuous mode. The MS^2^ spectra were generated in data dependent (Full MS/dd-MS^2^) scan mode with a resolution of 17 500 FWHM. Data acquisition was obtained with Xcalibur™ Software 2.0 (Thermo Fisher Scientific, USA).

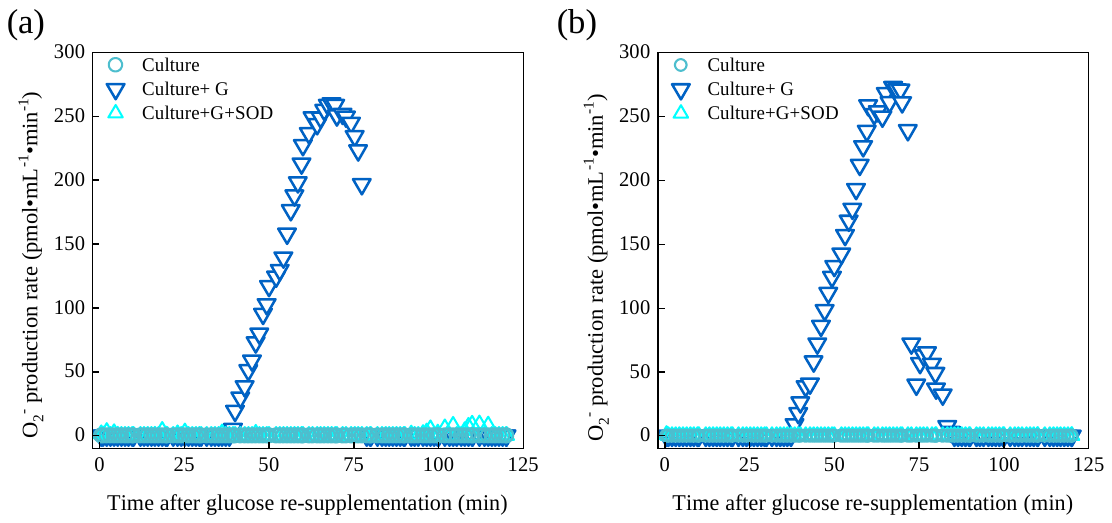

**Supplementary Fig. 1** Superoxide CL signal intensity in 48-h *A.* QXT-31 cultures re-supplemented with/without sterile glucose. (a) and (b) are results of other biological replicates of Fig. 1a. G: Glucose.
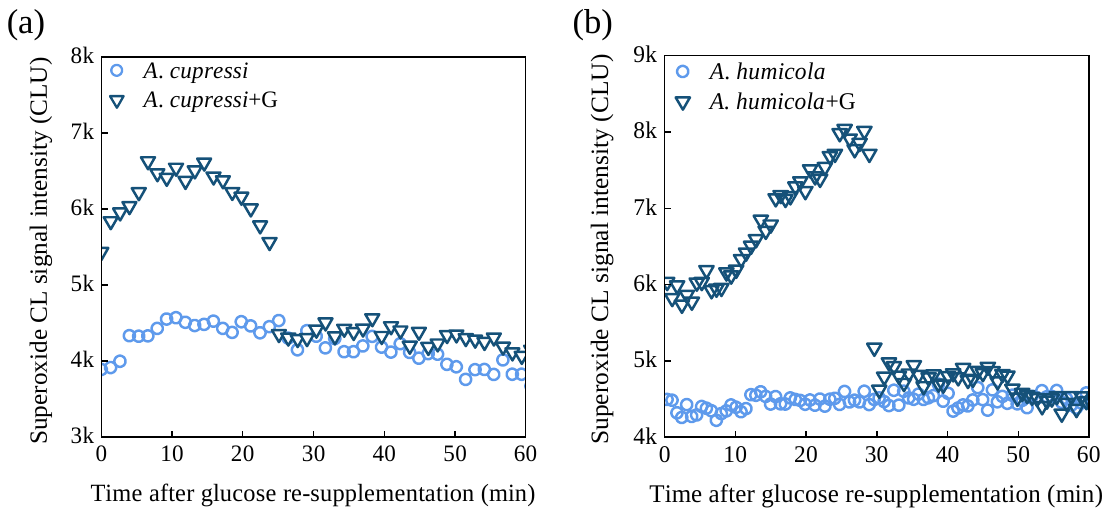

**Supplementary Fig. 2** Superoxide detection in cultures of two *Arthrobacter* strains after glucose re-supplementation. After glucose re-supplementation (50 mg•L^-1^), superoxide CL signal intensity was determined in (a) 4-d *A*. *cupressi* culture and (b) 24-h *A. humicola* culture. Both strains were grown in mPYG medium. G: Glucose.

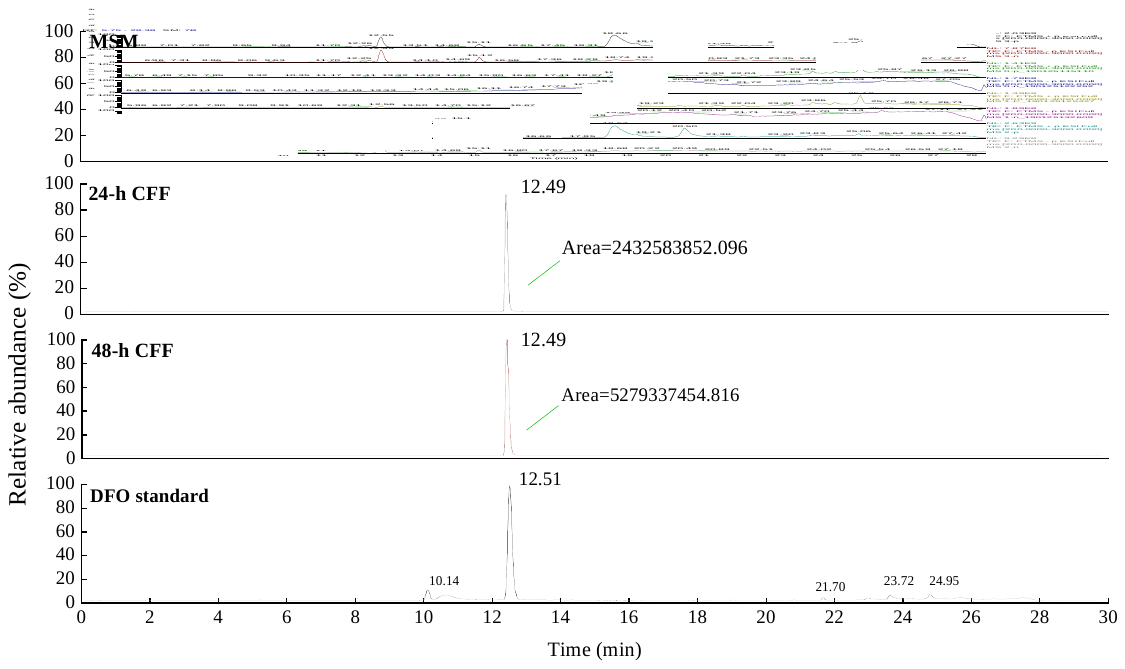

**Supplementary Fig. 3** Chromatograms of MSM, <3k Da CFF fractions in 24- and 48-h *A*. QXT-31cultures and 15.23 μM of DFO standard.

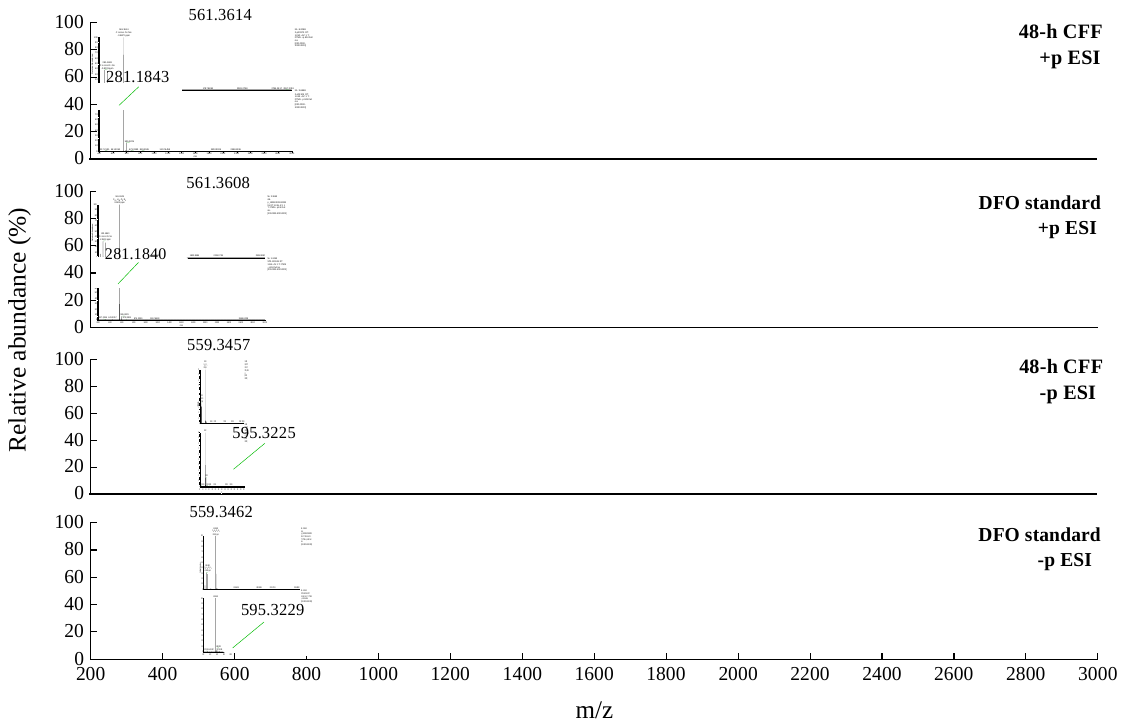

**Supplementary Fig. 4** Comparison of primary electrospray ionization mass spectrogram between suspected chromatographic peak of <3 kDa CFF fraction in 48-h *A*. QXT-31 culture and DFO standard (15.23 μM) under positive (+p) and negative (-p) ionization modes.

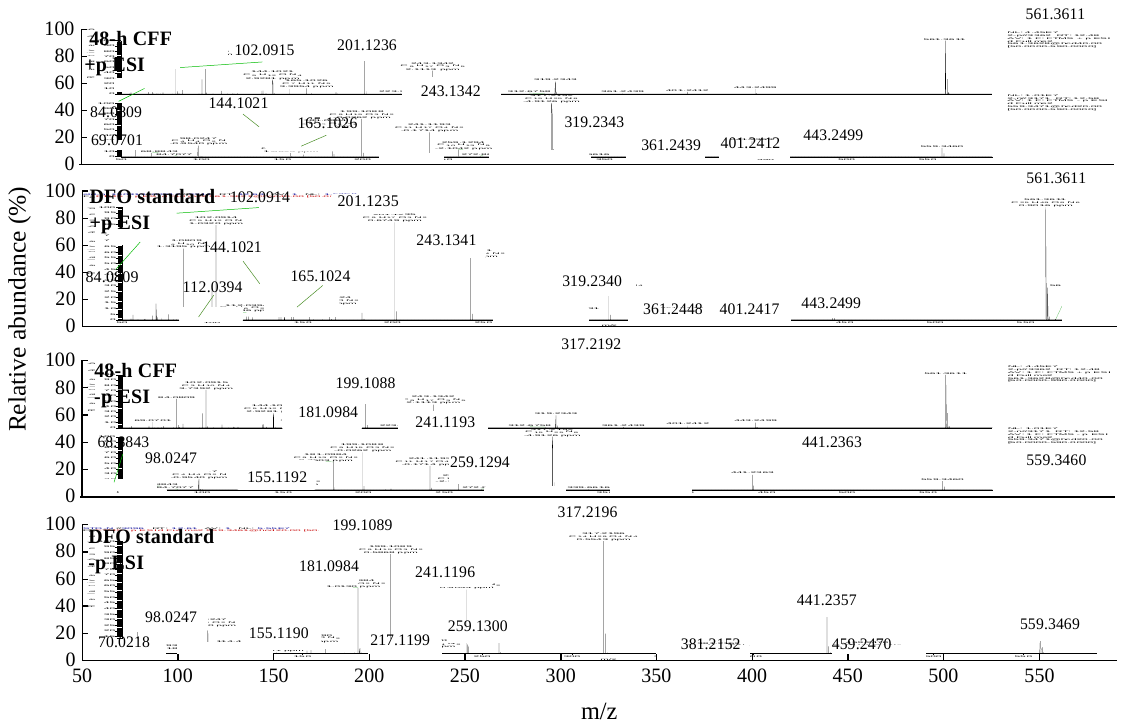

**Supplementary Fig. 5** Comparison of electrospray ionization secondary mass spectrogram between suspected chromatographic peak of <3 kDa CFF fraction in 48-h *A*. QXT-31 culture and DFO standard (15.23 μM) under positive (+p) and negative (-p) ionization modes.

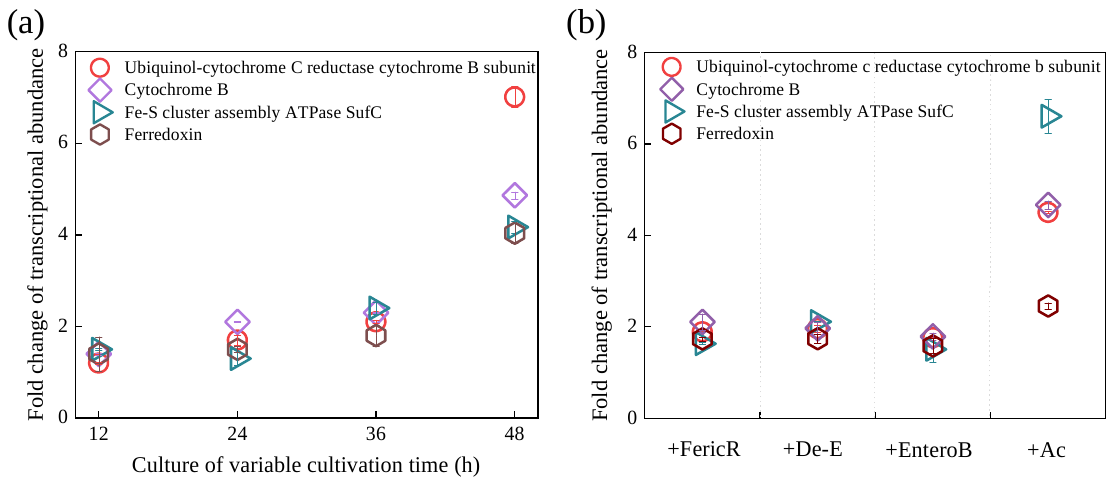

**Supplementary Fig. 6** Siderophores induced transcriptional up-regulation of several genes encoding iron-related and iron-bearing proteins. (a) Changes in the transcriptional abundance of genes encoding iron-related and iron-bearing proteins after ~2 μM sterile DFO was added to *A.* QXT-31 cultures. After DFO addition, cells in 0.5 mL of culture were harvested by centrifugation (10 000 *g*, 4 °C, 3 min) at 0.5 h, 1 h, 1.5 h, and 2 h, with four cell samples collected at four time points then mixed well as an RNA-Seq sample. (b) Changes in the transcription level of genes encoding iron-related and iron-bearing proteins in 36-h culture of *A.* QXT-31 after supplementation (2 μM) of acetomenadione acid (Ac), deferrioxamine E (De-E), enterobactin (EnteroB), and ferrichrome (FericR). RNA-Seq samples were prepared according to that of DFO. Data are means ± average deviation of two biological replicates.

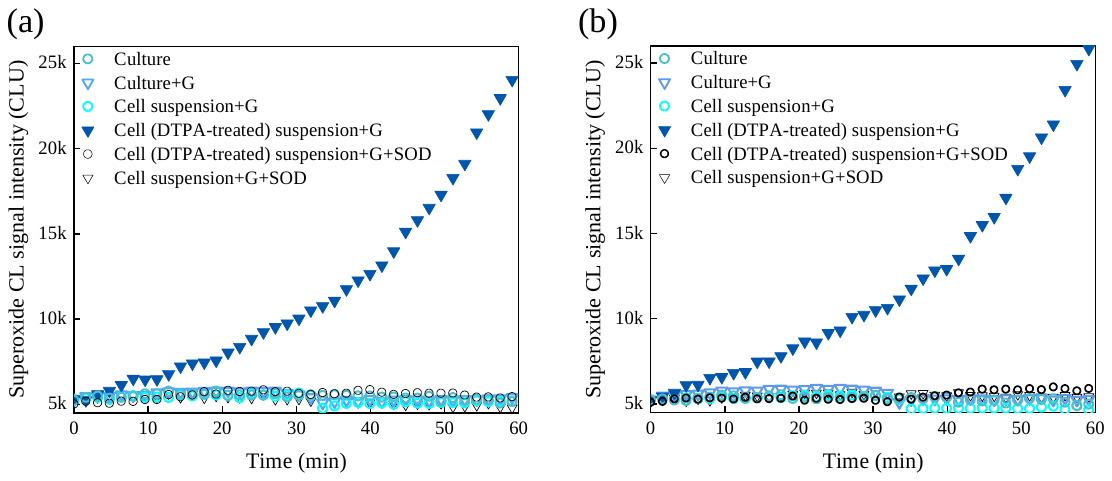

**Supplementary Fig. 7** Superoxide CL signal intensity in glucose-re-supplemented 24-h *A*. QXT-31 culture and suspension of DTPA-treated 24-h *A*. QXT-31 cells. (a) and (b) are results of other biological replicates of Fig. 4b. G: Glucose.

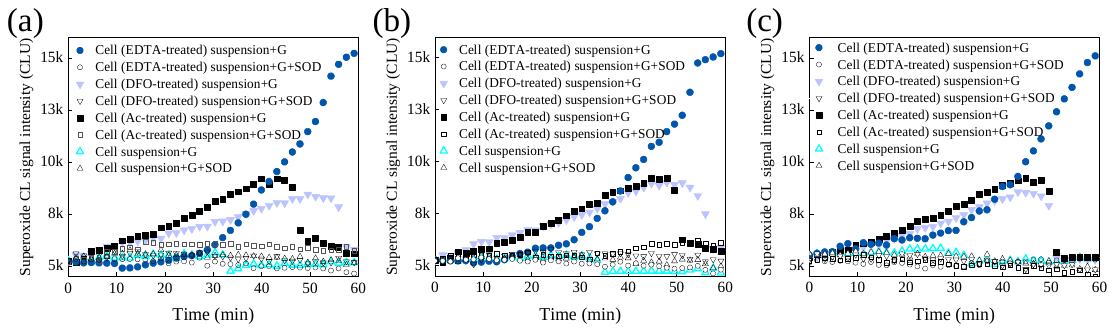

**Supplementary Fig. 8** Siderophore provoked *A*. QXT-31 cells to produce superoxide. Superoxide CL signal intensity in glucose-re-supplemented (50 mg•L^-1^) 24-h *A*. QXT-31 cultures and suspensions (in CFF of 24-h culture) of siderophore-treated 24-h *A*. QXT-31 cells. 24-h *A*. QXT-31 culture was supplemented with/without 10 μM sterile siderophore (DFO/EDTA/acetohydroxamic acid (Ac)), followed by shaking at 170 rpm at 30 °C in an oscillating incubator for 10 min. Cells were collected by centrifugation (718 *g*, 30 °C, 10 min), and then suspended in CFF of 24-h culture (siderophores free). SOD (120 kU•L^-1^) was added to generate superoxide-free controls. G: Glucose.

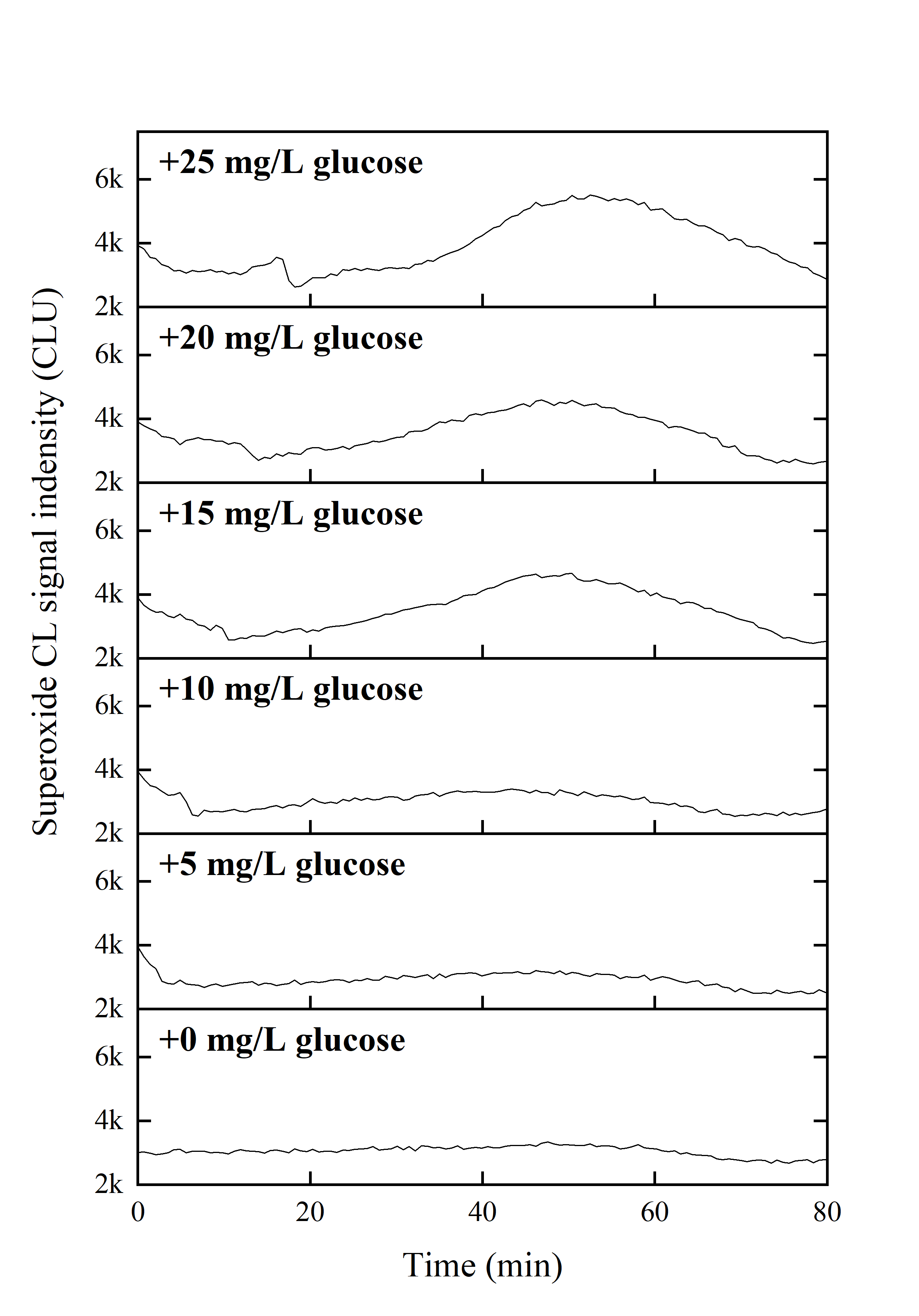
**Supplementary Fig. 9** Superoxide CL signal intensity in 48-h *A*. QXT-31 culture after glucose supplementation at various concentrations. Superoxide was monitored on a microplate reader after 180 μL of culture was added to each prepared microplate well (reagents added in advance).
